# Serum glycoprotein biomarker validation for esophageal adenocarcinoma and application to Barrett’s surveillance

**DOI:** 10.1101/281220

**Authors:** Alok K. Shah, Gunter Hartel, Ian Brown, Clay Winterford, Renhua Na, Kim-Anh Lê Cao, Bradley A. Spicer, Michelle Dunstone, Wayne A. Phillips, Reginald V. Lord, Andrew P. Barbour, David I. Watson, Virendra Joshi, David C. Whiteman, Michelle M. Hill

**Author notes:** **Correspondence:** Michelle M. Hill, PhD, QIMR Berghofer Medical Research Institute, Locked Bag 2000, Royal Brisbane Hospital, 300 Herston Road, Herston QLD 4029 Australia.

## Abstract

**BACKGROUND & AIMS:** Esophageal adenocarcinoma (EAC) is thought to develop from asymptomatic Barrett’s esophagus (BE) with a low annual rate of conversion. Current endoscopy surveillance for BE patients is probably not cost-effective. Previously, we discovered serum glycoprotein biomarker candidates which could discriminate BE patients from EAC. Here, we aimed to validate candidate serum glycoprotein biomarkers in independent cohorts, and to develop a biomarker panel for BE surveillance.

**METHODS:** Serum glycoprotein biomarker candidates were measured in 301 serum samples collected from Australia (4 states) and USA (1 clinic) using lectin magnetic bead array (LeMBA) coupled multiple reaction monitoring mass spectrometry (MRM-MS). The area under receiver operating characteristic curve was calculated as a measure of discrimination, and multivariate recursive partitioning was used to formulate a multi-marker panel for BE surveillance.

**RESULTS:** Different glycoforms of complement C9 (C9), gelsolin (GSN), serum paraoxonase/arylesterase 1 (PON1) and serum paraoxonase/lactonase 3 (PON3) were validated as diagnostic glycoprotein biomarker candidates for EAC across both cohorts. A panel of 10 serum glycoproteins accurately discriminated BE patients not requiring intervention [BE+/-low grade dysplasia] from those requiring intervention [BE with high grade dysplasia (BE-HGD) or EAC]. Tissue expression of C9 was found to be induced in BE, dysplastic BE and EAC. In longitudinal samples from subjects that have progressed towards EAC, levels of serum C9 glycoforms were increased with disease progression.

**CONCLUSIONS:** Further prospective clinical validation of the confirmed biomarker candidates in a large cohort is warranted. A first-line BE surveillance blood test may be developed based on these findings.

**Abbreviations:**

AAL: *Aleuria aurantia* lectin
%CV: % Co-efficient of variation
AUROC: Area under receiver operating characteristics curve
BE: Barrett’s esophagus
BE-HGD: Barrett’s esophagus with high-grade dysplasia
BE-ID: Barrett’s esophagus which is indefinite for dysplasia
BE-LGD: Barrett’s esophagus with low-grade dysplasia
BMI: Body mass index
C1QB: Complement C1q subcomponent subunit B
C2: Complement C2
C3: Complement C3
C4B: Complement C4-B
C4BPA: C4b-binding protein alpha chain
C4BPB: C4b-binding protein beta chain
C9: Complement component C9
CFB: Complement factor B
CFI: Complement factor I
CI: Confidence interval
CP: Ceruloplasmin
EAC: Esophageal adenocarcinoma
EPHA: Erythroagglutinin from *Phaseolus vulgaris*
FFPE: Formalin-fixed, paraffin-embedded
GERD: Gastroesophageal reflux disease
GSN: Gelsolin
JAC: Jacalin from *Artocarpus integrifolia*
LeMBA: Lectin magnetic bead array
MRM-MS: Multiple reaction monitoring-mass spectrometry
NPL: *Narcissus pseudonarcissus* lectin
NSE: Non-specialized epithelium
OR: Odds ratio
PGLYRP2: N-acetylmuramoyl-L-alanine amidase
PON1: Serum paraoxonase/arylesterase 1
PON3: Serum paraoxonase/lactonase 3
RBP4: Retinol-binding protein 4
SERPINA4: Kallistatin
SIS: Stable isotope-labeled internal standard

## INTRODUCTION

Esophageal cancer is the sixth most common cause of cancer related mortality in men, with 3-fold higher rates in men than women (1, 2). Of the two main histological subtypes (adenocarcinoma and squamous cell carcinoma), the incidence of esophageal adenocarcinoma (EAC) has been rising continuously in Western countries, and accounts for the majority of cases (3–5). Despite aggressive treatment, EAC has a 5-year survival of less than 20% (6). EAC is thought to develop from the metaplastic condition Barrett’s esophagus (BE) as a consequence of gastroesophageal reflux disease (GERD) through a metaplasia–dysplasia–adenocarcinoma sequence (Figure 1A) (7–9).

**Figure 1.**
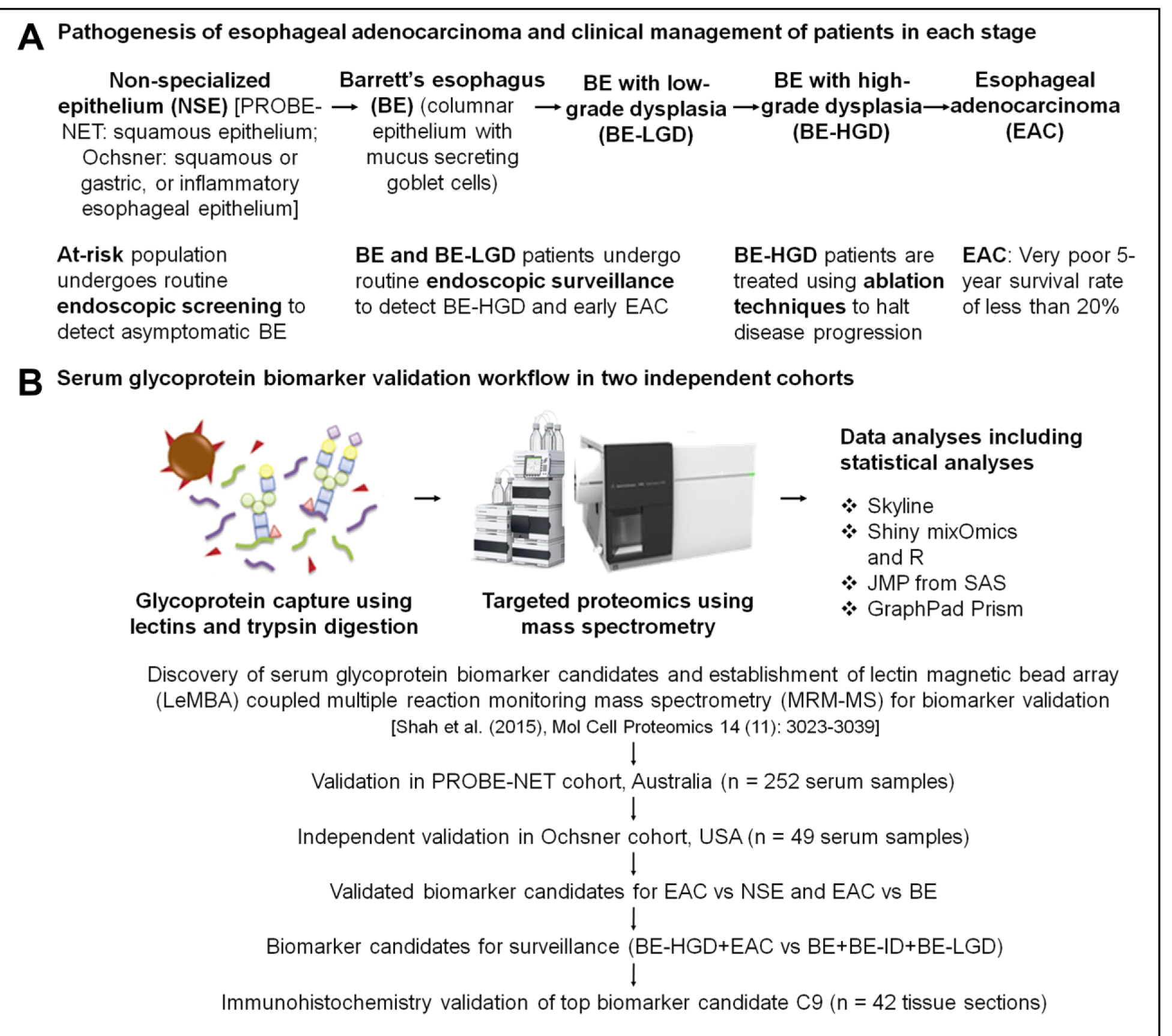
**(A)** Pathogenesis of esophageal adenocarcinoma (**EAC**) and clinical management of patients during each stage. In response to exposure to gastric and bile acid, non-specialized esophageal epithelium (**NSE**) converts to Barrett’s esophagus (**BE**), and may progress through low grade dysplasia (**LGD**) and high grade dysplasia (**HGD**) stages to EAC. Patients at high risk of BE undergo endoscopic screening to detect asymptomatic metaplastic BE condition. Patients with BE or BE-LGD undergo endoscopy-biopsy surveillance to detect BE-HGD for endoscopic treatments. **(B)** Workflow of the study. A total of 301 serum samples from two different patient cohorts were subjected to lectin magnetic bead array-coupled multiple reaction monitoring mass spectrometry (LeMBA-MRM-MS). Biomarker candidates for EAC and surveillance were identified by statistical analysis. Tissue expression of top glycoprotein biomarker candidate complement C9 was evaluated by immunohistochemistry.

Currently, BE patients usually undergo endoscopy-biopsy surveillance with the degree of dysplasia assessed by histopathology as a biomarker to monitor risk of neoplastic progression (10). Patients diagnosed with high grade dysplasia (BE-HGD) are treated with endoscopic mucosal resection, radiofrequency ablation or surgery, in an attempt to halt further disease progression (10–12). The significant cost of endoscopy plus the low annual progression rate to HGD or EAC means that the cost-effectiveness of endoscopic surveillance is questioned at the population level (13–16). Furthermore the evaluation of dysplasia in tissue biopsies by histopathology is prone to inter-observer variability and sampling error (17). A less costly and minimally invasive diagnostic procedure is needed for cost-effective screening and surveillance of at-risk populations (18, 19).

As the first step to developing blood-based EAC diagnostic test, we focused on differential glycosylation of serum glycoproteins during EAC pathogenesis. We established a new glycoprotein biomarker pipeline which couples lectin-based glycoprotein isolation with state-of-the-art discovery and targeted proteomics (20–23). We then applied it to identify and verify changes in lectin binding profile of serum glycoproteins between healthy, BE and EAC patients (22). Here, we report results from validation in independent cohorts, and evaluation of biomarker panels for surveillance of BE patients.

## EXPERIMENTAL PROCEDURES

### Clinical cohorts

Ethical approval was obtained from all participating institutions, and all patients provided informed consent to participate in the studies. We investigated two independent cohorts recruited in Australia and the United States of America, respectively. The Australian samples were selected from participants recruited into The Progression of Barrett’s Esophagus to Cancer Network (PROBE-NET) study across 4 states (New South Wales, Queensland, South Australia and Victoria). A total of 252 serum samples collected from 242 patients were analyzed [Normal – 43, BE – 65, BE-LGD – 39, BE-HGD – 35, and EAC – 60, at baseline]. Of the 252 serum samples, 10 samples were from patients who progressed to the subsequent stage of the disease while 242 were baseline serum.

Independently, a cross-sectional cohort of 49 serum samples collected at Ochsner Health System, New Orleans, United States were also analyzed [Normal – 14, BE – 13, Barrett’s mucosa which is indefinite for dysplasia (BE-ID) – 3, BE-LGD – 2, BE-HGD – 7, and EAC – 10]. Some patients in the Normal group had a history of Barrett’s related pathology and received endoscopic treatments. These patients were confirmed to have no Barrett’s mucosa by histology at the time of serum collection, hence the normal groups will be called non-specialized epithelium [NSE, epithelium without Barrett’s mucosa].

The clinical diagnosis linked to the sample was based on histological examination of biopsies taken at the same endoscopy. The diagnoses were provided to the researcher performing the biomarker measurements to allow batch randomization design for the assay. Serum samples were stored at -80°C, aliquoted, and shipped to the Translational Research Institute, Brisbane on dry ice for this study. Formalin-fixed, paraffin-embedded (FFPE) tissue sections from the Ochsner Health System were selected and shipped to Brisbane, Australia for immunohistochemistry.

Both PROBE-NET and the Ochsner cohort provided information on patients’ age, sex and body mass index (BMI, calculated as weight (kg) / [height (m)]^2^); whereas ethnicity was provided by the Ochsner cohort only and education, alcohol drinking and tobacco smoking was only available in the PROBE-NET cohort. Data on demographics and lifestyle factors were compared among different clinical and histological groups using Pearson’s chi-square test or Fisher’s exact test as appropriate. *P* < 0.05 was considered to be statistically significant. Analyses were performed using SAS 9.4 software.

### Serum glycoprotein biomarker measurement and analysis

Serum glycoprotein biomarker candidates were measured using our previously reported lectin magnetic bead array (20, 21)-coupled multiple reaction monitoring (MRM) mass spectrometry assay (24). In this method, lectin binding is used to isolate glycoproteins with particular glycan structures. Based on our previous work for BE/EAC (22, 23), four lectins were selected for the independent validation cohorts, namely AAL, EPHA, JAC, and NPL (Vector Laboratories, Burlingame, CA). PROBE-NET and Ochsner cohorts were independently analyzed with block randomization design for each cohort. The PROBE-NET method measured 365 peptides belonging to 106 proteins while Ochsner method measured 381 peptides belonging to 115 proteins, inclusive of standard peptides. Detailed methods are provided in Supplemental Methods and Supplemental Table 1.

**Table 1.**
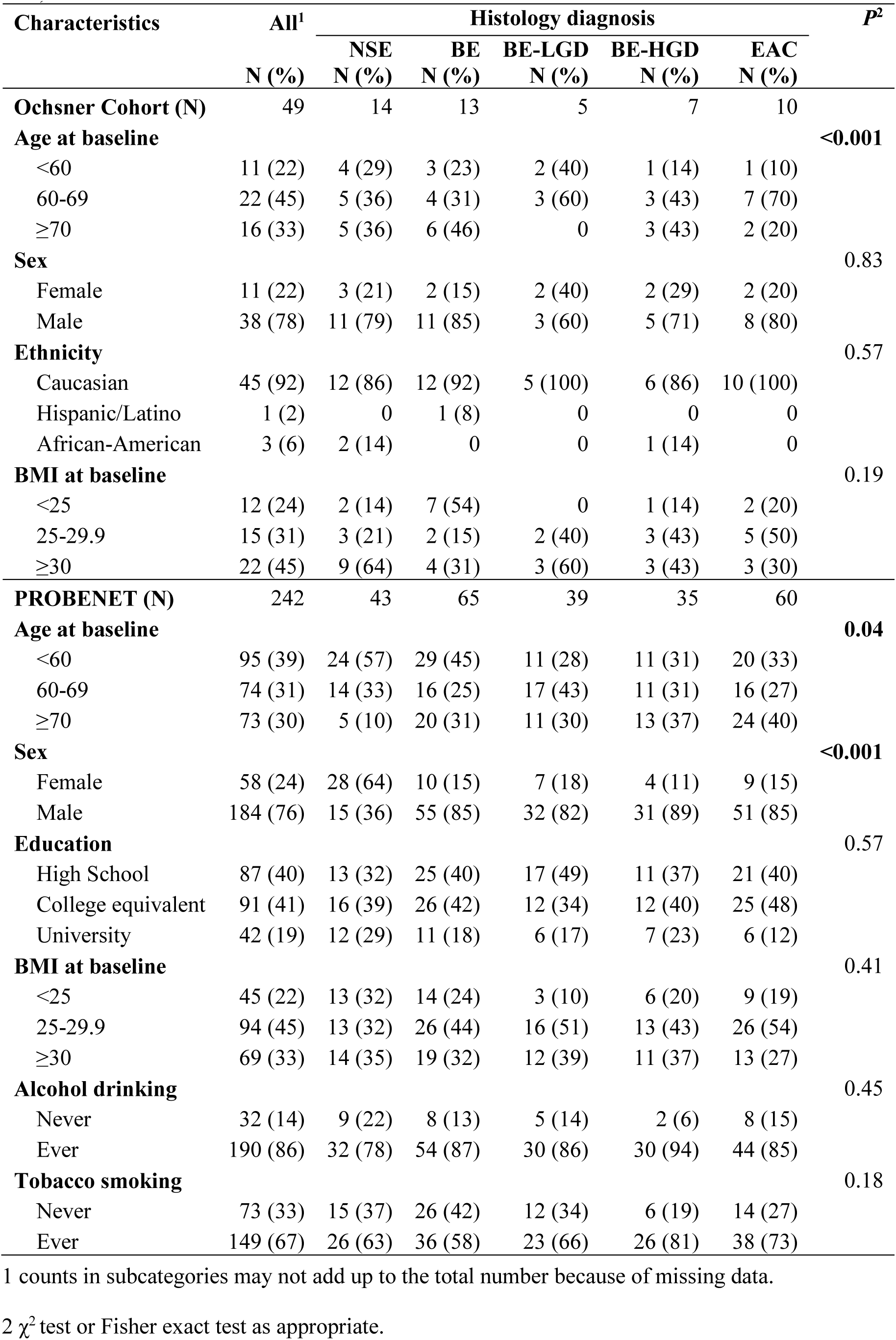
Baseline demographics, histologic and clinical characteristics of the study cohorts (N = <u>291).</u>

| Characteristics | All<sup>1</sup><br>N (%) | Histology diagnosis |  |  |  |  | P<sup>2</sup> |
| --- | --- | --- | --- | --- | --- | --- | --- |
|  |  | NSE<br>N (%) | BE<br>N (%) | BE-LGD<br>N (%) | BE-HGD<br>N (%) | EAC<br>N (%) |  |
| **Ochsner Cohort (N)** | 49 | 14 | 13 | 5 | 7 | 10 |  |
| **Age at baseline** |  |  |  |  |  |  | **<0.001** |
| <60 | 11 (22) | 4 (29) | 3 (23) | 2 (40) | 1 (14) | 1 (10) |  |
| 60-69 | 22 (45) | 5 (36) | 4 (31) | 3 (60) | 3 (43) | 7 (70) |  |
| ≥70 | 16 (33) | 5 (36) | 6 (46) | 0 | 3 (43) | 2 (20) |  |
| **Sex** |  |  |  |  |  |  | 0.83 |
| Female | 11 (22) | 3 (21) | 2 (15) | 2 (40) | 2 (29) | 2 (20) |  |
| Male | 38 (78) | 11 (79) | 11 (85) | 3 (60) | 5 (71) | 8 (80) |  |
| **Ethnicity** |  |  |  |  |  |  | 0.57 |
| Caucasian | 45 (92) | 12 (86) | 12 (92) | 5 (100) | 6 (86) | 10 (100) |  |
| Hispanic/Latino | 1 (2) | 0 | 1 (8) | 0 | 0 | 0 |  |
| African-American | 3 (6) | 2 (14) | 0 | 0 | 1 (14) | 0 |  |
| **BMI at baseline** |  |  |  |  |  |  | 0.19 |
| <25 | 12 (24) | 2 (14) | 7 (54) | 0 | 1 (14) | 2 (20) |  |
| 25-29.9 | 15 (31) | 3 (21) | 2 (15) | 2 (40) | 3 (43) | 5 (50) |  |
| ≥30 | 22 (45) | 9 (64) | 4 (31) | 3 (60) | 3 (43) | 3 (30) |  |
| **PROBENET (N)** | 242 | 43 | 65 | 39 | 35 | 60 |  |
| **Age at baseline** |  |  |  |  |  |  | **0.04** |
| <60 | 95 (39) | 24 (57) | 29 (45) | 11 (28) | 11 (31) | 20 (33) |  |
| 60-69 | 74 (31) | 14 (33) | 16 (25) | 17 (43) | 11 (31) | 16 (27) |  |
| ≥70 | 73 (30) | 5 (10) | 20 (31) | 11 (30) | 13 (37) | 24 (40) |  |
| **Sex** |  |  |  |  |  |  | **<0.001** |
| Female | 58 (24) | 28 (64) | 10 (15) | 7 (18) | 4 (11) | 9 (15) |  |
| Male | 184 (76) | 15 (36) | 55 (85) | 32 (82) | 31 (89) | 51 (85) |  |
| **Education** |  |  |  |  |  |  | 0.57 |
| High School | 87 (40) | 13 (32) | 25 (40) | 17 (49) | 11 (37) | 21 (40) |  |
| College equivalent | 91 (41) | 16 (39) | 26 (42) | 12 (34) | 12 (40) | 25 (48) |  |
| University | 42 (19) | 12 (29) | 11 (18) | 6 (17) | 7 (23) | 6 (12) |  |
| **BMI at baseline** |  |  |  |  |  |  | 0.41 |
| <25 | 45 (22) | 13 (32) | 14 (24) | 3 (10) | 6 (20) | 9 (19) |  |
| 25-29.9 | 94 (45) | 13 (32) | 26 (44) | 16 (51) | 13 (43) | 26 (54) |  |
| ≥30 | 69 (33) | 14 (35) | 19 (32) | 12 (39) | 11 (37) | 13 (27) |  |
| **Alcohol drinking** |  |  |  |  |  |  | 0.45 |
| Never | 32 (14) | 9 (22) | 8 (13) | 5 (14) | 2 (6) | 8 (15) |  |
| Ever | 190 (86) | 32 (78) | 54 (87) | 30 (86) | 30 (94) | 44 (85) |  |
| **Tobacco smoking** |  |  |  |  |  |  | 0.18 |
| Never | 73 (33) | 15 (37) | 26 (42) | 12 (34) | 6 (19) | 14 (27) |  |
| Ever | 149 (67) | 26 (63) | 36 (58) | 23 (66) | 26 (81) | 38 (73) |  |

1 counts in subcategories may not add up to the total number because of missing data.
2 $\chi^2$ test or Fisher exact test as appropriate.

Skyline was used for inspecting and processing MRM data (25). The quality of acquired datasets (Supplemental Table 2A for PROBE-NET cohort and Supplemental Table 2B for Ochsner cohort) were evaluated by % coefficient of variation (% CV) of spiked-in stable isotope-labeled internal standard (SIS) peptides and peptides derived from the spiked-in internal standard protein chicken ovalbumin. The data were normalized using the median intensity of 8 SIS peptides. Peptide intensities were converted into protein intensity with Pearson correlation coefficient cut-off set at 0.6 (22). As recently highlighted by others (26), this step serves as quality control for peptide level measurements resulting in a robust protein level quantitative dataset for down-stream statistical analysis. The normalized protein intensities were transformed using the natural logarithm and z-scores were calculated for downstream statistical analysis.

**Table 2.** Glycoprotein biomarker panel performance for BE surveillance (BE-HGD+EAC vs BE+BE-ID+BE-LGD) in PROBE-NET and Ochsner cohorts.

| Multimarker panel | PROBE-NET cohort |  |  | Ochsner cohort |  |
| --- | --- | --- | --- | --- | --- |
|  | AUROC | Sensitivity | Specificity | Sensitivity | Specificity |
| **Panel 1:**<br>C9_EPHA + PON1_AAL +<br>SERPINA4_EPHA +<br>RBP4_AAL + C1QB_JAC +<br>PGLYRP2_NPL | 0.83 | 83% | 69% | 61% | 65% |
| **Panel 2:** Panel 1 +<br>CP_NPL + C4BPB_JAC +<br>C2_EPHA + C4B_AAL | 0.93 | 86% | 85% | 61% | 71% |

AAL = *Aleuria aurantia* lectin; AUROC = area under receiver operating characteristic curve; BE = Barrett's esophagus; BE-HGD = BE with high-grade dysplasia; BE-LGD = BE with low-grade dysplasia; C1QB = Complement C1q subcomponent subunit B; C2 = Complement C2; C4B = Complement C4-B; C4BPB = C4b-binding protein beta chain; C9 = complement component C9; CP = Ceruloplasmin; EAC = esophageal adenocarcinoma; EPHA = erythroagglutinin *Phaseolus vulgaris*; GSN = gelsolin; JAC = jacalin from *Artocarpus integrifolia*; NPL = *Narcissus pseudonarcissus* lectin; PGLYRP2 = N-acetylmuramoyl-L-alanine amidase; PON1 = serum paraoxonase/arylesterase 1; PON3 = serum paraoxonase/lactonase 3; RBP4 = Retinol-binding protein 4; SERPINA4 = Kallistatin.

JMP Pro 13.2 (SAS Institute, Inc., Cary, NC, USA) was used for univariate and multivariate biomarker statistical analyses. Univariate logistic regressions were conducted on EAC vs NSE outcome, EAC vs BE outcome, and the surveillance outcome (BE-HGD or EAC vs BE or BE-ID or BE-LGD), against each of the glycoprotein_lectin biomarker candidates. Odds ratios (ORs) with 95% Wald confidence intervals (95% CIs), area under receiver operating characteristic curves (AUROCs) and Likelihood Ratio *P* values were calculated for both PROBE-NET and Ochsner cohorts. Recursive partitioning (also known as Classification and Regression Trees, CART) was used to identify a multivariate panel of markers that would discriminate between surveillance outcomes (BE-HGD+EAC vs BE+BE-ID+BE-LGD). The PROBE-NET dataset was used as the training set to develop predictive models, and the Ochsner dataset was used as the validation set. The set of 217 markers that were available in both PROBE-NET and Ochsner datasets was used in the training set, as well as baseline characteristics, including age, gender, and BMI. To avoid overfitting, models were limited to 6, 8, and 10 biomarkers. The prediction formulas derived from PROBE-NET were then applied to the Ochsner dataset to determine sensitivity and specificity in the validation set.

### Immunohistochemical analysis

Immunohistochemical staining was performed on 42 tissue sections collected from 34 patients using anti-C9 primary antibody (Sigma Aldrich #SAB4503059). Detailed immunohistochemistry protocol is described in Supplemental Methods. To confirm the staining specificity, the anti-C9 antibody was neutralized with C9 recombinant protein before application to the slide, as reported in Supplemental Methods and Supplemental Figure 1. Staining intensities in squamous mucosa, columnar epithelium without intestinal metaplasia, Barrett’s esophagus mucosa (with intestinal metaplasia), dysplasia (low and high grade), EAC and inflammatory infiltrate were scored by a specialist gastrointestinal pathologist. Each component, if present in the tissue, was scored separately using a 4 grade assessment of intensity (0 no staining, 1+ weak staining, 2+ moderate staining, 3+ strong staining). Where the staining was non-uniform in a component, the maximum intensity of staining was used for the score, providing at least 10% of the cells of that component stained to this intensity. Due to limited numbers, for statistical analysis, we combined dysplasia with EAC group; similarly, those with 3+ staining in Barrett’s mucosa were combined with 2+ staining group. The relationship between histological features and the staining intensity was evaluated using Fisher exact test.

## RESULTS

Workflow of this study is depicted in Figure 1B. Table 1 details the baseline demographic and clinical characteristics of the Australian (PROBE-NET) and Ochsner cohorts.

Univariate analysis of the PROBE-NET cohort dataset revealed 46 and 54 biomarker candidates with *P*<0.05 for EAC vs NSE and EAC vs BE comparisons respectively (Supplemental Table 3). Data for the top 10 biomarkers that differentiate EAC from NSE, and EAC from BE in PROBE-NET cohort are shown in Figure 2A and 2B, respectively. Out of these candidates, 16 candidates for EAC vs NSE comparison and 9 candidates for EAC vs BE comparison were also significantly different in the Ochsner cohort, confirming these glycoproteins as validated biomarkers. As illustrated in Figure 2C, 8 validated biomarkers overlap between the two lists. These biomarkers are potentially most useful, being able to distinguish EAC from NSE and BE. Interestingly, the 8 biomarkers comprised of different glycosylated forms of 4 proteins, namely, complement C9 (C9), gelsolin (GSN), serum paraoxonase/arylesterase 1 (PON1), serum paraoxonase/lactonase 3 (PON3).

**Figure 2.**
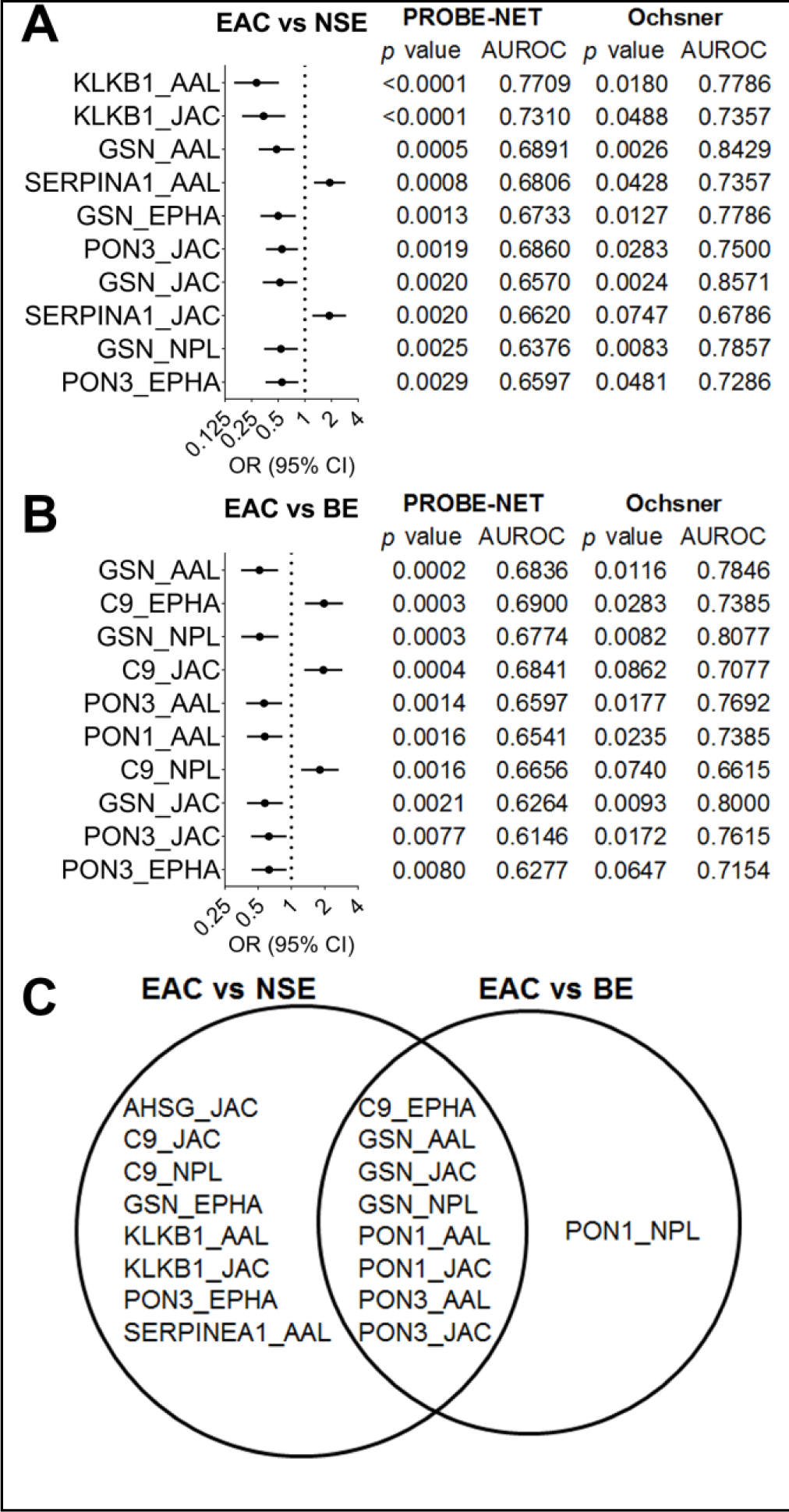
Serum glycoprotein biomarker validation in PROBE-NET and Ochsner cohorts. Top 10 glycoprotein_lectin biomarkers that differentiate **(A)** esophageal adenocarcinoma (EAC) from non-specialized epithelium (NSE) and **(B)** EAC from Barrett’s esophagus (BE) in PROBE-NET and Ochsner cohorts. **(C)** Overlap of validated biomarkers (*P*<0.05). Biomarkers are shown by gene name and lectin affinity, and ordered by *P* value. Horizontal bar indicates odds ratio (OR) with 95% Wald confidence intervals. Likelihood Ratio *P* values were calculated to test statistical significance in the univariate logistic regressions. Area under receiver operating characteristic curve (AUROC) indicates diagnostic ability of individual biomarker candidate. AAL = *Aleuria aurantia* lectin; C9 = complement C9; EPHA = erythroagglutinin *Phaseolus vulgaris*; GSN = gelsolin; JAC = jacalin from *Artocarpus integrifolia*; KLKB1 = plasma kallikrein; NPL = *Narcissus pseudonarcissus* lectin; PON1 = serum paraoxonase/arylesterase 1; PON3 = serum paraoxonase/lactonase 3; SERPINA1 = alpha-1-antitrypsin.

Next we considered BMI as a potential confounding factor for EAC biomarker validation. Correlation analysis between BMI and biomarker levels in all PROBE-NET samples revealed no substantial correlation (|r|<0.6) (Supplemental Table 4), although three proteins namely complement C3 (C3), C4b-binding protein alpha chain (C4BPA), and complement factor I (CFI) with multiple lectin pull-downs showed positive correlations with BMI in NSE patients only (|r| 0.6234-0.7069). Importantly, none of the top 10 biomarkers showed strong correlation with BMI.

### Biomarkers for BE surveillance

Having confirmed univariate biomarkers for detection of EAC from NSE and BE in independent cohorts, we next evaluated the ability of serum glycoproteins to be used as a surveillance tool for BE and BE-LGD patients, i.e. to distinguish between patients who require treatment (BE-HGD and EAC) and those who do not (BE, BE-ID and BE-LGD). Eight biomarkers that showed AUROC > 0.6 in a BE surveillance setting for both PROBE-NET and Ochsner cohort (Figure 3, Supplemental Table 3), comprising of 3 glycoforms of GSN, 2 glycoforms of C9, AAL-binding PON1, AAL-binding PON3, as well as EPHA-binding, Complement factor B (CFB).

**Figure 3.**
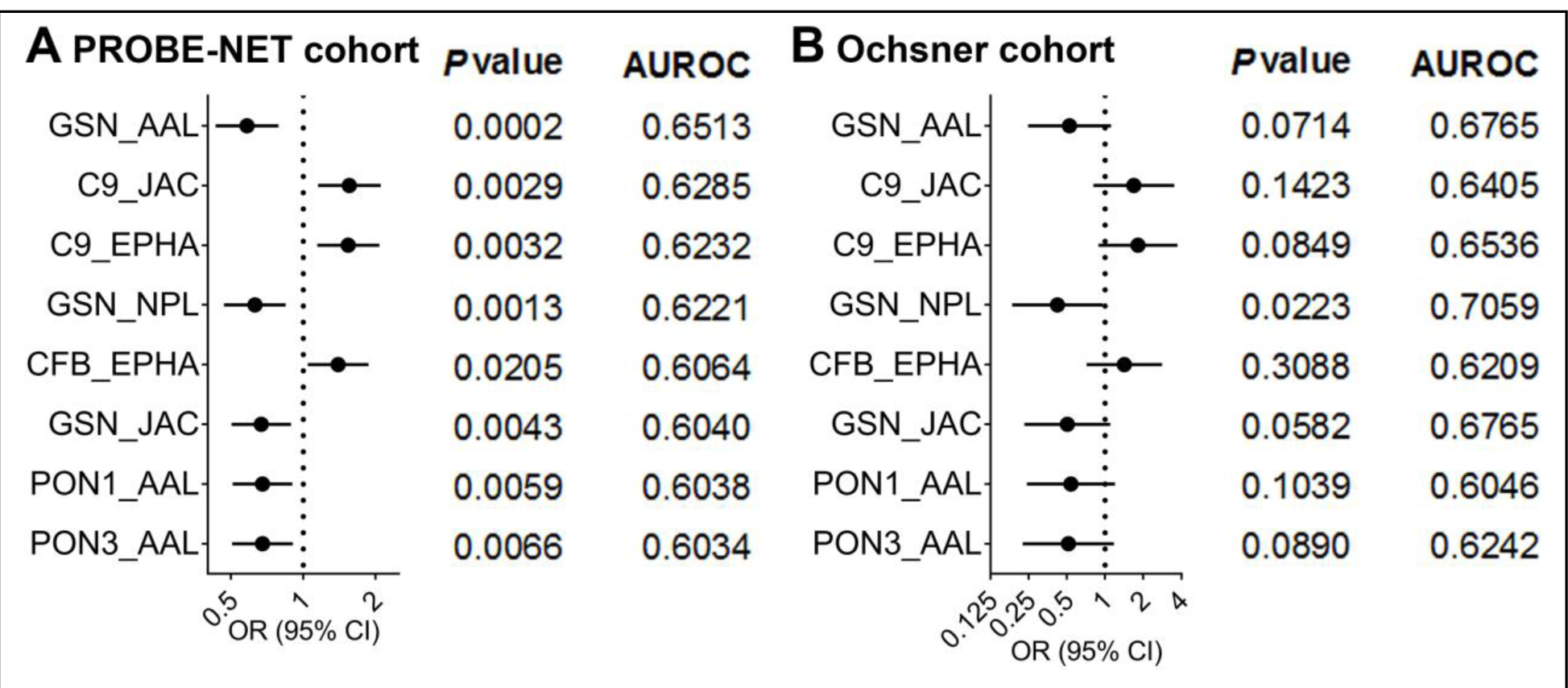
Serum glycoprotein biomarkers for BE surveillance in PROBE-NET and Ochsner cohorts. Glycoproteins that distinguish patients who require treatment (BE-HGD+EAC) from those who do not require treatment (BE+BE-ID+BE-LGD) with an area under receiver operating characteristic curve (AUROC) value of > 0.6 in both PROBE-NET and Ochsner cohorts are shown. Odds ratio (OR) with 95% Wald confidence intervals, likelihood ratio *P* values and AUROC values for **(A)** PROBE-NET cohort and **(B)** Ochsner cohort. AAL = *Aleuria aurantia* lectin; AUROC = area under receiver operating characteristic curve; BE = Barrett’s esophagus; BE-HGD = BE with high-grade dysplasia; BE-LGD = BE with low-grade dysplasia; C9 = complement C9; CFB = complement factor B; CI = confidence interval; EAC = esophageal adenocarcinoma; EPHA = erythroagglutinin *Phaseolus vulgaris*; GSN = gelsolin; JAC = jacalin from *Artocarpus integrifolia*; NPL = *Narcissus pseudonarcissus* lectin; PON1 = serum paraoxonase/arylesterase 1; PON3 = serum paraoxonase/lactonase 3.

Next we sought to generate a multimarker panel for BE surveillance, using the PROBE-NET cohort for modeling and Ochsner cohort for model validation. The minimal panel of six biomarkers showed 0.83 AUROC, 83% specificity and 67% sensitivity for the PROBE-NET cohort, and a moderate specificity of 61% and sensitivity of 63% for the Ochsner cohort (Table 2). Addition of 4 more biomarkers to the panel increased the AUROC to 0.93, and improved the specificity and sensitivity measures for PROBE-NET, as well as the sensitivity for Ochsner cohort (Table 2).

### Complement C9 expression in BE and EAC tissue

In agreement with our previous finding of complement pathway dysregulation in EAC pathogenesis (22), 5 of the 10 glycoprotein biomarker candidates in the final surveillance biomarker panel (Table 2) belonged to the complement pathway. As a first step to evaluate alterations of the complement pathway in EAC at a tissue level, we optimized immunohistochemistry staining for the top candidate complement C9. Staining specificity of the method was confirmed by neutralization of the antibody with recombinant C9 protein prior to staining (Supplemental Figure 1). We then evaluated expression of C9 in esophageal tissue sections from a subset of the Ochsner cohort. As shown in Figure 4A, C9 was detected in BE and EAC, but not squamous esophageal epithelium. Dysplastic BE showed particularly strong staining in the plasma membrane and/or cytoplasm (Figure 4A). Strong staining in immune infiltrates served as an expected positive control. Quantitation of staining intensity score against the tissue phenotype (Figure 4B) showed statistically significant association between C9 expression levels and histology groups of squamous epithelium, columnar epithelium, Barrett’s mucosa, dysplasia/EAC (*P*<0.001).

**Figure 4.**
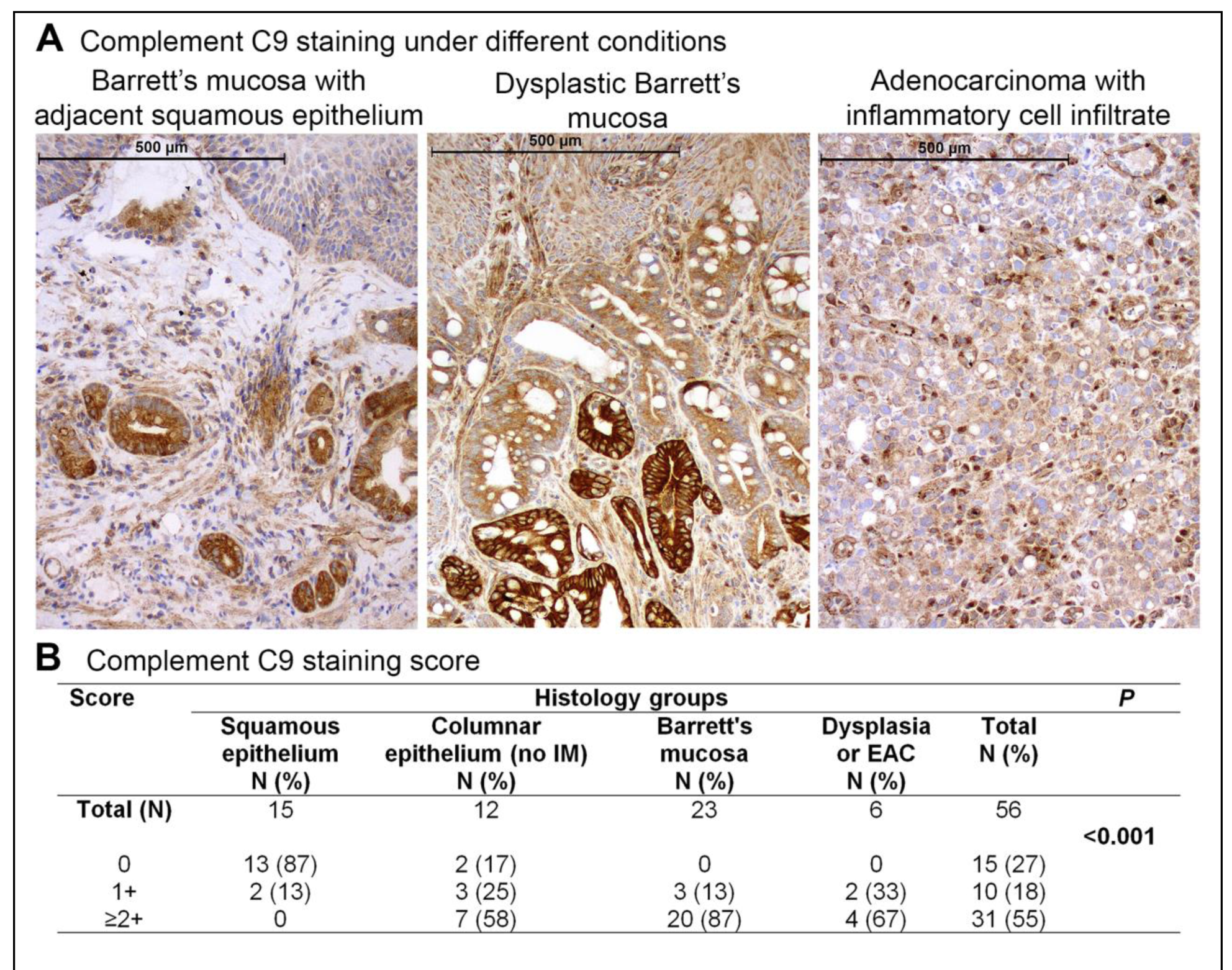
Immunohistochemical analysis of complement C9 (C9) in lower esophageal biopsies. **(A)** A panel of representative images of C9 staining in squamous epithelium, Barrett’s mucosa, dysplastic epithelium, and esophageal adenocarcinoma conditions. The left image shows 2+ staining in the non-dysplastic Barrett’s epithelium. The middle image shows 1+ and 3+ staining in the BE. The right image shows 2+ staining in cancer cells and 3+ staining in inflammatory cells. **(B)** Summary of C9 staining intensity according to histology groups. *P* value from Fisher exact test. EAC = esophageal adenocarcinoma; IM = intestinal metaplasia.

### Serum complement C9 glycoforms in progressor samples

As an additional evaluation, we examined C9 lectin pull-down levels in samples from PROBE-NET participants who had progressed from BE to BE-LGD (N = 4), BE-LGD to BE-HGD (N = 3), BE to BE-HGD (N = 1), BE to EAC (N = 1), and NSE to gastric type mucosa (N = 1) phenotype during follow-up. The progression time varied from 110 days up to 5 years. Significant elevation of C9_EPHA and C9_NPL was observed following progression in this small patient cohort (Figure 5).

**Figure 5.**
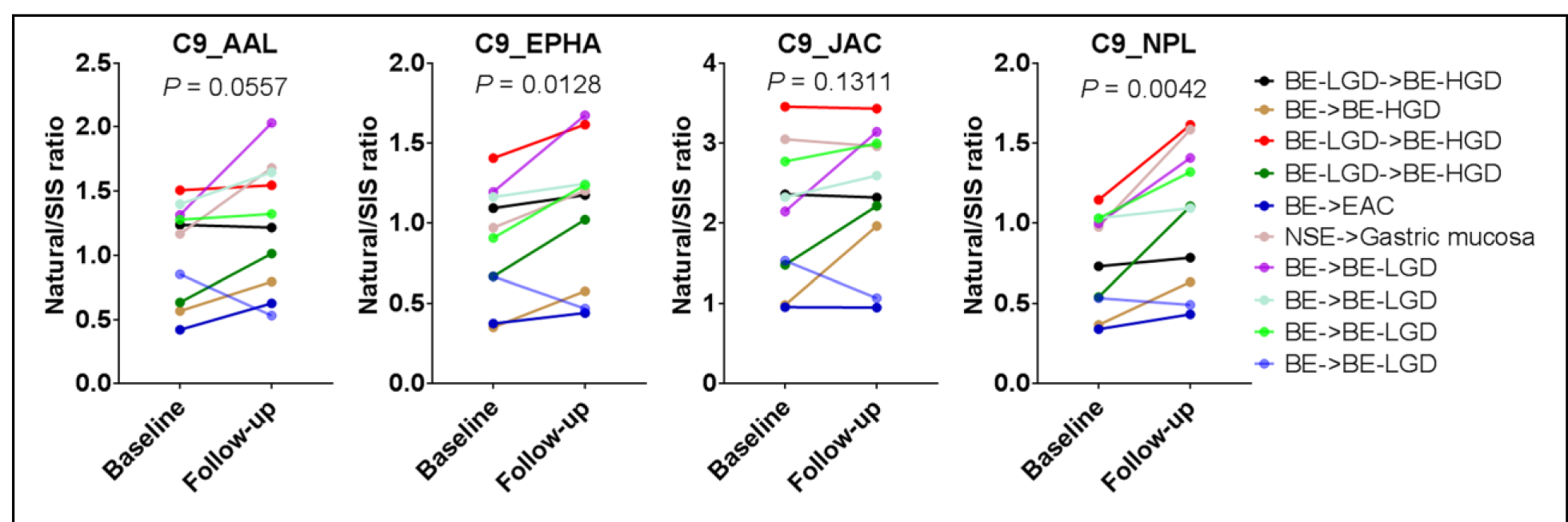
Serum complement C9 (C9) levels in specific lectin-pulldowns in PROBE-NET progressor serum samples. C9 levels in lectin pull-downs are expressed as ratio of intensity of natural peptide LSPIYNLVPVK to stable isotope labeled spiked-in peptide response. The changes in C9 in lectin pull-down samples of participants who progressed from one stage (baseline) to another stage of the disease at follow-up. *P* value from paired t-test. AAL = *Aleuria aurantia* lectin; BE = Barrett’s esophagus; BE-HGD = BE with high-grade dysplasia; BE-LGD = BE with low-grade dysplasia; C9 = complement C9; EAC = esophageal adenocarcinoma; EPHA = erythroagglutinin *Phaseolus vulgaris*; JAC = jacalin from *Artocarpus integrifolia*; NPL = *Narcissus pseudonarcissus* lectin; SIS = stable isotope standard.

## DISCUSSION

This study advances our previous serum glycoprotein research (22, 23) and provides a critical breakthrough towards developing cost-effective EAC surveillance. Over the years, several studies have been carried out to identify circulatory biomarker candidates to diagnose BE-dysplasia-EAC disease spectrum (19, 27). These studies have explored genetic alterations in cell free circulating DNA (28), serum miRNA changes (29), circulatory tumor cells (30), glycan profile alteration in serum (31, 32), circulatory autoantibodies (against cancer antigens) (33), volatile organic compounds found in breath analysis (34), metabolic changes in urine (35), and a panel of serum proteins (36) as promising diagnostic biomarker candidates for BE and/or EAC. However, none of these biomarker candidates have progressed to from bench to bedside, likely due to the lack of subsequent validation studies in large independent cohort of patients. Here, we have addressed this gap by validating eight serum glycoprotein biomarkers for EAC in two independent patient cohorts including dysplastic samples.

In addition to demonstrating the robustness of our mass spectrometry based glycoprotein-centric proteomics workflow for biomarker validation, the current study confirmed our previous finding of complement activation in EAC (22). The complement system consists of a cascade of circulating proteases that are locally activated leading to the formation of the membrane attack complex on the immunogen and recruitment of phagocytes. Complement components are predominantly expressed and secreted into the plasma by the liver but are also found to be expressed in other tissues (37), but some complement proteins may be expressed by tumor cells (38). While complement components are primarily involved in mediating innate immune response, recent studies have revealed an apparently paradoxical tumor-promoting role of the complement system (38, 39). Complement C3 has been reported to play an autocrine role in ovarian and lung cancer tumor growth (40). C5a is elevated in serum of lung cancer patients (41) and increases the invasiveness of C5aR+ tumors (42). Fucosylated C9 was previously reported to be elevated in the serum of lung cancer patients (43).

Two previous publications reported on complement component changes in BE and EAC. Bobryshev et al. reported reduced C1q expression in dendritic cells and macrophages in the epithelium of BE and EAC, and suggest this to be an immune-escape mechanism (44). Song et al. identified complement pathway proteins, complement C3 and complement C1r subcomponent to be increased in serum collected from BE-HGD and EAC patients respectively as compared to disease free individuals (45). The current study is the first to report C9 protein expression in BE and EAC cells, as demonstrated on tissue sections. In addition to validating our previous finding of elevated circulating C9 glycoforms in EAC, we further confirmed the utility of serum C9_JAC and C9_EPHA for BE surveillance in independent cohorts (Figure 3). The latter result suggests a pathological change in the secretion, glycosylation, or expression of C9 during the progression of BE to BE-HGD/EAC. Modulation of C9 expression is unlikely to be the mechanism, since we detected strong C9 staining in both BE and BE-HGD tissue. Interestingly, bile (deoxycholic acid) treatment was reported to altered glycosylation and Golgi structure in esophageal epithelial and BE cells, resulting in impaired protein secretion via the classical pathway (46). Hence, C9 may be sequentially regulated by expression level and glycosylation/secretion during BE-EAC progression. Further studies are required to determine the precise molecular mechanisms.

The strengths of this study include the validation of biomarkers in independent cohorts, evaluation of biomarker panel for BE surveillance, and the use of immunohistochemistry to determine the cellular origin of the top serum glycoprotein biomarker. There are several limitations to our study. While our glycoprotein biomarker pipeline allows high-throughput glycoprotein biomarker discovery, the glycosylation site and glycan structural changes are not determined at early stages. Furthermore, the functional consequences of these glycosylation changes in cancer progression also remain to be evaluated. The small number of progressor samples available is a limitation, not only for the current study, but for evaluation of BE/EAC biomarkers in general. International collaborations and longitudinal cohort sampling are therefore required to advance the search for EAC biomarkers. Finally, the tissue expression of C9 and other candidates also need to be confirmed in a larger number of samples.

In summary, we have validated a number of serum glycoprotein biomarkers that warrant further clinical testing in large independent cohorts including longitudinal patient samples. Further development of these biomarkers to a blood test can aid the current endoscopy-biopsy surveillance program of BE patients for early detection of dysplastic progression.

## Author contributions

MMH conceived and supervised the project. AKS designed and performed the proteomics experiments including data analysis. AKS, GH, RN, and KALC undertook statistical analysis. WAP, RVL, APB, DIW, DCW, and VJ contributed to sample collection. RN, DCW, and VJ selected the patient samples for the study. CW developed, optimized and performed the immunohistochemistry. IB evaluated and scored immunohistochemistry slides. BAS and MD contributed to recombinant C9 protein expression and purification. AKS, GH and MMH wrote the manuscript. DCW helped with critical revision of the manuscript. All authors edited and approved the manuscript.

We thank Dr Thomas Hennessey (Agilent Technologies) and Mr Elliot McElroy (Agilent Technologies) for technical assistance, and Mr Cris Molina (Ochsner) for clinical research management support.

### Grant support

This work was supported by The University of Queensland-Ochsner Seed Fund for Collaborative Research Grant 2014, The University of Queensland Faculty of Medicine and Biomedical Science Cancer Bequest Grant 2014, and internal funding support from The University of Queensland Diamantina Institute 2015. PROBE-NET was supported by a National Health and Medical Research Council (NHMRC) Centre of Research Excellence grant (APP1040947). KALC is a recipient of NHMRC Career Development Fellowship (APP1087415). DCW is supported by an NHMRC Research Fellowship (APP1058522). The sponsors had no influence on the study design, collection, analysis, and interpretation of data.

### Disclosures

The authors have no conflicts of interest to disclose.

